# Anisotropic fluorescence emission by diatoms modifies the underwater oceanic light field

**DOI:** 10.1101/2025.05.28.656578

**Authors:** Joan Salvador Font-Muñoz, Jorge Arrieta, Marc Sourisseau, Idan Tuval, Gotzon Basterretxea

## Abstract

Fluorescence in phytoplankton and other autotrophic organisms originates within the cell chloroplasts, where a fraction of the absorbed solar radiation is reemitted at longer wavelengths by photopigments. While traditionally employed as an indicator of physiological status, emerging evidence suggests that natural chlorophyll fluorescence (ChlF) may also play unanticipated functional roles in the marine environment. Here, we examine the ChlF emission fields generated by pennate planktonic diatoms, a key phytoplankton group playing a critical role in global biogeochemical cycles. Using cell micromanipulation experiments, we demonstrate that the ChlF emitted by *Pseudonitzschia fraudulenta* (Bacillariophyceae) is markedly anisotropic, a feature attributed to both the cells’ elongated morphology and the arrangement of chloroplasts within the cytoplasm. In these diatoms, fluorescence is preferentially emitted in the transapical direction, accounting for up to 35% of total emission. However, peak ChlF intensities occur at the cell apices, suggesting that the silica frustule focuses fluorescent light emission along the longitudinal axis. At elevated cell densities (∼10^6^ cells/L), the underwater light field is modulated by the combined effects of ChlF emission anisotropy and the preferential alignment of diatom cells within the water column. Numerical simulations indicate that ChlF intensity can vary by up to 15% depending on whether cells are predominantly aligned —commonly in stratified water columns— or randomly oriented. These results suggest that diatom-driven modulation of the light field through structured ChlF emission may influence microscale optical environments, with potential consequences for processes ranging from intercellular signaling to large-scale phytoplankton dynamics, including remote sensing–based assessments of phytoplankton physiology.

**SIGNIFICANCE STATEMENT:** Phytoplankton are microscopic organisms that underpin marine food webs and play a central role in global carbon cycling. This study shows that certain diatoms, a major group of phytoplankton, emit chlorophyll fluorescence in a strongly directional manner, influenced by their shape, internal structure, and collective orientation. At natural population densities, this anisotropic fluorescence emission can significantly modify the distribution of light underwater. Such biologically driven alterations to the ocean light field may impact phytoplankton interactions, growth dynamics, and how they are observed by satellite remote sensing. These findings uncover a previously overlooked mechanism by which phytoplankton influence their optical environment, with important implications for understanding marine ecosystem processes, improving interpretation of ocean color data, and enhancing models of ocean productivity.

## INTRODUCTION

The optical phenomenon of fluorescence in which short-wavelength light is absorbed and re-emitted at longer wavelengths, also referred to as biofluorescence or autofluoresce, is widespread in the natural world. This trait is found in diverse life forms, from microorganisms to mammals, reptiles, and birds, inhabiting a wide range of ecosystems, both marine and terrestrial that contain biomolecules, namely endogenous fluorophores, that emit fluorescence when exposed to light (1–5). More than 300 different natural biomolecules, including chlorophyll, phycobiliproteins, porphyrins, chitin, or green-fluorescent protein (GFP)-like fluorescent proteins have been described to produce fluorescence (6–8).

Fluorescent biomolecules present chemical properties in terms of conjugated double bonds, aromatic or more complex structures with oxidized and crosslinked bonds, ensuring an energy status able to permit electronic transitions matching with the energy of light in the UV-visible-near-IR spectral range (9). The fundamental principles of fluorescence photophysics are governed by the molecular transitions that occur upon absorption of a photon by a fluorophore, which can occupy multiple discrete energy states, each corresponding to a distinct configuration of its molecular orbitals. Upon photon absorption, the fluorophore is excited from its base level or ground state to a higher electronic energy state, followed by rapid non-radiative relaxation to the lowest vibrational level of the excited state, from which it returns to the ground state via the emission of a photon (10).

In terrestrial ecosystems, the ambient light environment spans the full visible spectrum, which poses a significant challenge for detecting the relatively weak fluorescent emission due to its overlap with the broad and intense background illumination(11, 12). In contrast, marine ecosystems are ideal for organisms to perceive, without interference, the fluorescent light emitted by their conspecifics or other organisms (13). This is due to the specific optical properties of seawater, which progressively attenuate the overall intensity of sunlight and exercise a selective absorption of longer wavelengths in the visible spectrum. As other wavelengths are absorbed with increasing depths, the ocean becomes an environment dominated by short wavelengths, essentially a uniform blue light spectrum (470-490 nm; (14, 15). The conditions observed are ideal for fluorescence to emerge, as it relies on blue-UV light re-emission, and the incidence of other wavelengths is marginal. It is remarkable to observe how different life forms have leveraged these unique optical characteristics of seawater to exploit fluorescence for a multitude of functions, ranging from prey and symbiont attraction, photoprotection, and photoenhancement, mimicry, aposematism, and inter- and intraspecific communication (8). The favorable conditions in the sea for fluorescent-related functionalities could explain the prevalence of biofluorescence in marine organisms (see Lagorio *et al.*, 2015).

Sunlight is essential for photosynthesis and, therefore, plays a major role in determining the spatial distribution of marine phytoplankton, ultimately shaping life in the upper ocean. Photopigments such as chlorophyll-a (Chl-a), located within the chloroplasts of photosynthetic organisms, mediate the absorption of light and transfer the captured energy to the photosystems, where it drives the production of ATP and NADPH through the light-dependent reactions of photosynthesis. To avoid damage from light exceeding cell requirements, energy can be dissipated by Chl-a as heat or as red chlorophyll fluorescence (ChlF). Therefore, within each photosystem, energy is partitioned into three dynamic processes: the quantum yield of photochemistry, thermal energy dissipation, and re-emission of photons as fluorescence (17, 18). Consequently, the quantum yield of fluorescence (Φ_F_), defined as the ratio re-emitted photons to absorbed photons, has been linked to both Chl-a concentration and the photosynthetic rates of phytoplankton,, serving as a proxy for assessing physiological status and productivity. However, these relationships vary depending on several environmental and biological factors, including light history, nutritional status, photosynthetic capacity, physiological stress, photoprotective responses, and community composition (19–23).

Characteristically, ChlF emitted in cell chloroplasts is centered at around 683 nm (Q_y_ band) and has a half-maximal emission width of about 25 nm (Collins *et al.,* 1985; Kiefer *et al.,* 1989). It represents only a small fraction of all the excitation energy captured by the cell pigments (Φ_F_=2 to 5%; Walker, 1987; Falkowski & Raven, 2013). This red-light emission constitutes a strong and spectrally distinct signal that the ocean returns to space. Indeed, solar-induced fluorescence is the only signal detectable from space that can be unambiguously ascribed to life on Earth (26, 27).

While at first sight phytoplankton fluorescence could be considered a mere byproduct resulting from the deactivation pathway of photosynthesis, recent studies have proposed an unexpected role of fluorescence in congeneric cell-to-cell communication (3, 28, 29). In particular, certain functional groups, such as diatoms, may use ChlF for intercellular interactions, leading to synchronized population responses (28, 30). This mechanism of population-level coordination offers new insights into the social behavior of marine diatoms, which has traditionally been attributed primarily to chemical signaling via the exudation of specific information-bearing compounds known as infochemicals (31, 32). Nevertheless, while beaconing red light in the wave band of ChlF has been shown to trigger coordinated diatom responses, the characteristics and mechanisms of this signaling remain to be elucidated.

Here, through a series of laboratory experiments, we investigate the characteristics of the ChlF field emitted collectively and individually by pennate planktonic diatoms. Diatoms, unicellular eukaryotic algae belonging to the stramenopile lineage, rank among the most abundant photosynthetic organisms on Earth and play a pivotal role in primary production and nutrient cycling across both freshwater and marine ecosystems. They are ubiquitous and dominant in many coastal and oceanic regions, playing a major global trophic and biogeochemical role (33, 34). Diatoms are characterized by their unique silicified cell exoskeleton (frustule) comprising two valves (epitheca and hypotheca) and several girdle bands. Based on valve symmetry, diatoms are generally divided into the centric -with radial symmetry- and the pennate - with bilateral symmetry-, the latter group is further divided into the raphid and araphid diatoms, depending on the presence and absence of raphe, an opening in the silica cell wall from which mucilaginous material is secreted to facilitate cell motility.

We focus our study on the genus *Pseudo-nitzschia* (Heterokonta, Bacillariophyceae), a worldwide distributed marine planktonic raphid pennate diatom that includes several toxigenic species (35). The needle-like or obloid shell morphology of these diatoms can modify the propagation of light in the water column and the harvesting of light by cells (36–38). Through single-cell micromanipulation experiments, we measure the autofluorescence field emitted by individual pennate diatom cells and the characteristics emanating from the location of their chloroplasts within the cell structure. Then, we characterize the fluorescence field generated by a suspension of different species of pennate diatoms (*Pseudo-nitzschia fraudulenta, P. delicatissima,* and *P. pungens*) and its anisotropy variations attending to changes in population orientation. Building on cellular-level fluorescence measurements, we propose a population-level emission model and discuss the implications of the resulting anisotropic fluorescence field.

## RESULTS AND DISCUSSION

### Characterization of the single-cell ChlF field

It is widely recognized that the ecological success of diatoms in contemporary oceanic phytoplankton communities is largely attributable to their remarkable adaptability to fluctuating environmental conditions (see reviews by Mann, 1996; Fu *et al.*, 2022). Most raphid diatoms are mono- or diplastidic (39) and, in the case of *Pseudo-nitzschia*, two equally-sized chloroplasts positioned along the longitudinal axis of the cell, one on either side of the nucleus, and arranged symmetrically about the transapical plane of the cell (Fig. 1A). According to this chloroplast arrangement, ChlF emission is expected to be favored along the transapical axis (*x*-axis), where the frustule wall lies close to the chloroplasts and minimal internal cellular obstruction exists to attenuate the emitted red light (Figure 2). The cell was held by its valve face at the tip of a micropipette through gentle aspiration and rotated in place around its pervalvar axis at 0.01 rad intervals (Fig. 1B). The fluorescence at different rotation angles (0, π/4, π/2, 3π/4, π) is shown in Figure 2A. Red light is concentrated in the two emission foci centered in the chloroplasts. Total ChlF intensity - calculated as the sum of the intensity of each pixel - is consistently highest at π/2 and 3π/2, with minima at 0 and π.

**Figure 1.**
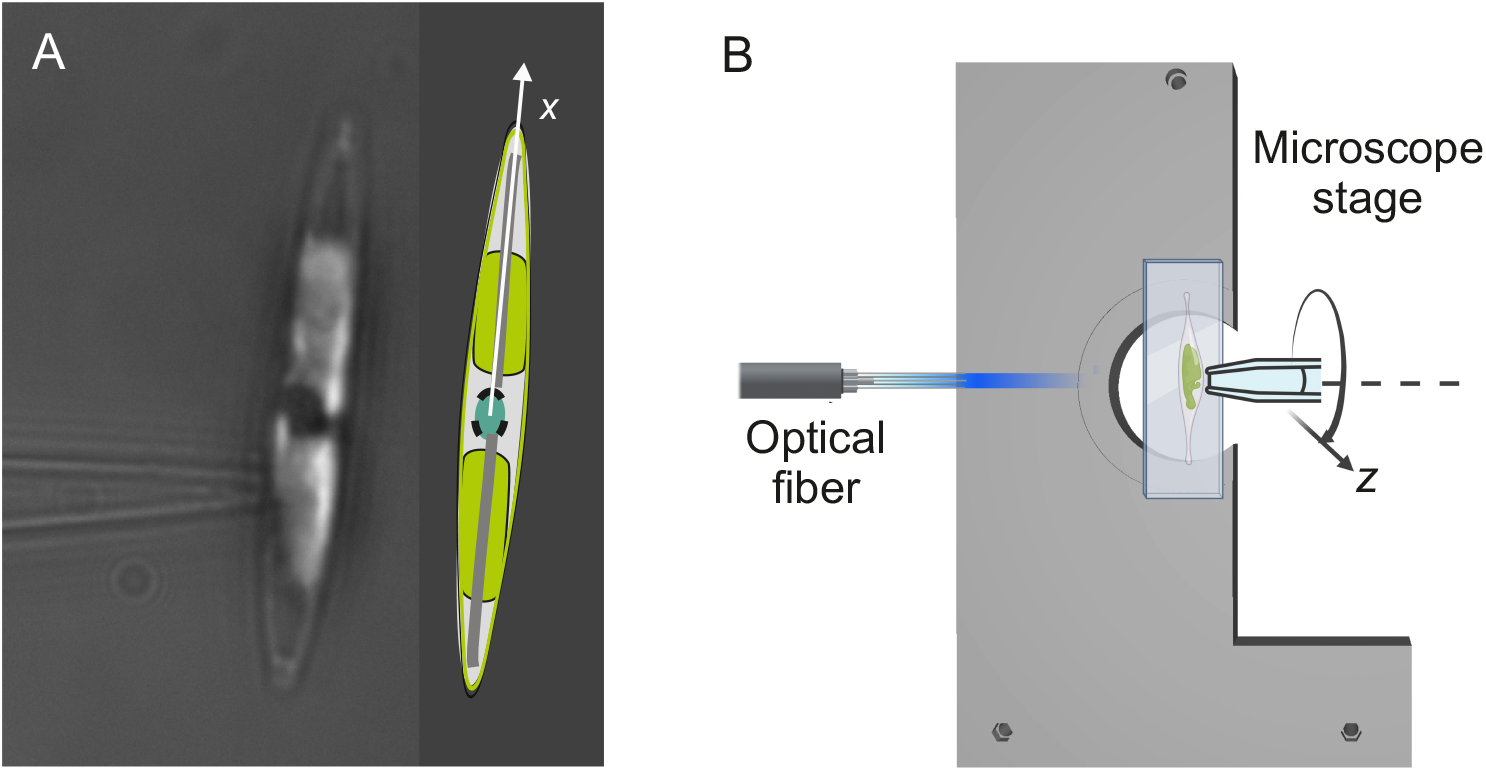
**(A)** *Pseudo-nitzschia fraudulenta* cell held by a micropipette and schematic showing the reference axis. **(B)** Experimental set-up allowing simultaneous fluorescence excitation and accurate single cell orientational positioning.

**Figure 2.**
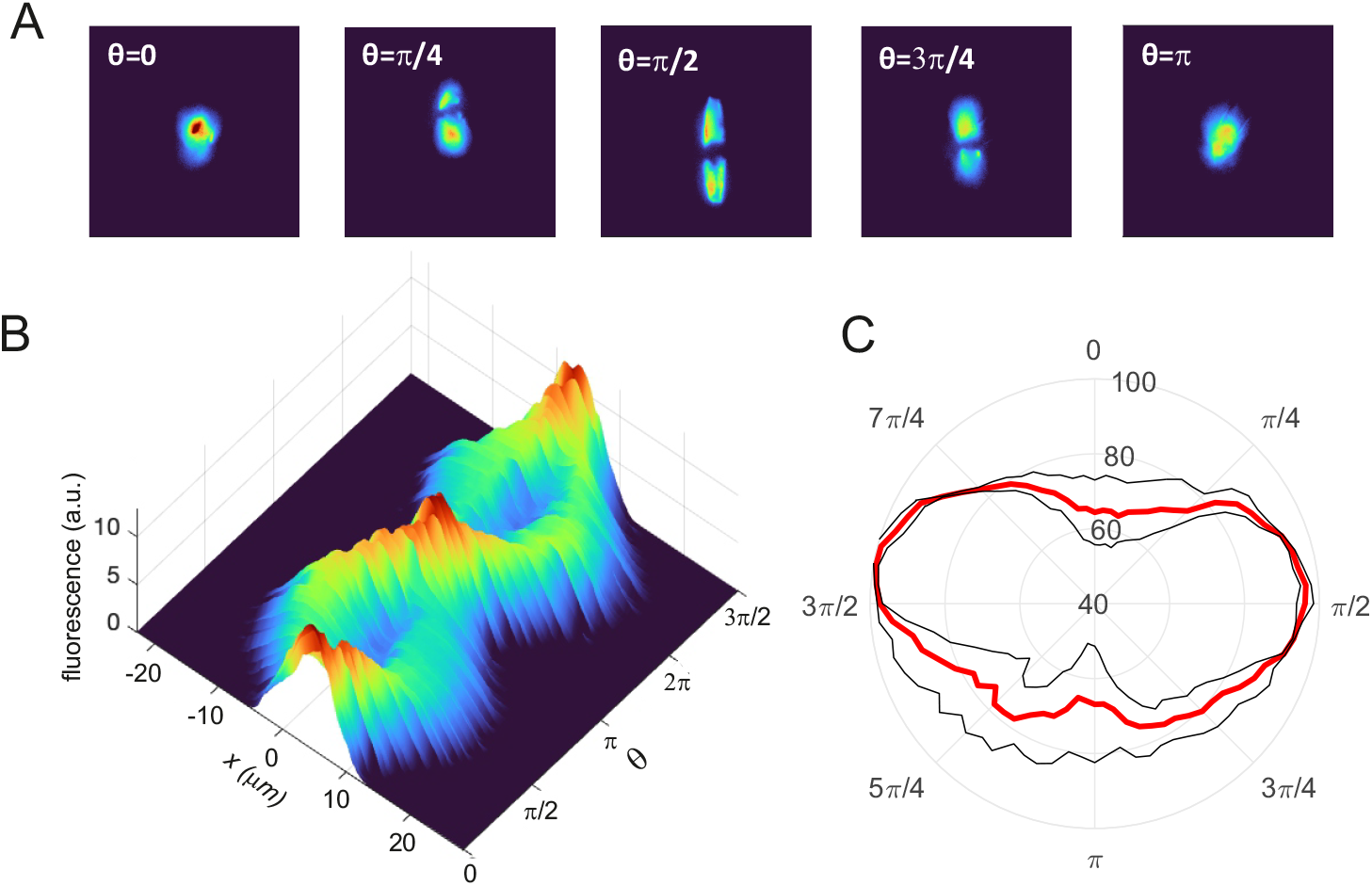
ChlF field emitted by a single *Pseudo-nitzschia fraudulenta* cell. **(A)** Images of the fluorescence emitted by a *Pseudo-nitzschia fraudulenta* cell at different angular orientations (*θ*). **(B)** Fluorescence intensity profiles of the cell (derived from the images) for each orientation, measured in steps of 0.02 rads. **(C)** Radial distribution of fluorescence at the single-cell level, depicting the total fluorescence intensity for each orientation (calculated as the sum of the intensities across all pixels in each image).

The anisotropy of the ChlF field can be described as the percentage variation between the maximum intensity in one direction and the minimum intensity in any other direction. This parameter is calculated as *r*=(I_max_−I_min_)/I_max_×100, where *r*=0% corresponds to an isotropic field. Based on this metric, the ChlF emitted at the cellular level shows a pronounced anisotropy, with r=35%. Also, a notch is observed in the fluorescence field in the range of ±0.35 rads from the apical axis (Fig. 2B). In this position, the autofluorescence from each of the two chloroplasts interferes with the nucleus and other internal structures, reducing ChlF. Strikingly, the highest fluorescence intensity values are approximately twice those measured at these alignment angles. These higher values are not explained by the proximity of the chloroplast to the cell wall since the cell periphery is richer in cytoplasmic content, and often numerous mitochondria are located in this apical position (41). More plausibly, it can be attributed to the optical properties of the frustules.

Owing to their periodic nanoporosity, diatom frustules exhibit remarkable optical properties, including the ability to guide and focus light along specific spatial orientations (42). According to Goessling *et al.* (2021), two main photobiological functions of the frustule have been proposed related to ambient light modulation: 1) modulation of photosynthetic light harvesting by wave-diffraction or forward scattering and 2) reduction of exposure to harmful radiation. However, although red light is transmitted more efficiently by the frustule than other wavelengths (44), knowledge about its role in transforming internally emitted red fluorescence remains limited. Only recently, using numerical simulations, D’Mello *et al.* (2022) reported the existence of light maxima across the apical axis for the pennate diatom *Nitzschia filiformis*. They suggested that the tapered end of the frustule functions as a butt-coupler, capable of funneling light in and out of the cell, which might enable light transfer between neighboring frustules in clusters. This indicates that optical interactions between cells could enable light sharing within clusters, similar to fiber-to-fiber coupling in optical systems.

### Anisotropy in the fluorescence field caused by preferential cell orientation

A collective characteristic of planktonic pennate diatoms emerges from their ability to orient themselves in response to the hydrodynamic conditions present in the water column. This behavior could influence the ambient fluorescence field generated by a diatom population. Under steady, low-shear conditions, elongated diatoms tend to align horizontally within the water column, frequently resulting in the formation of thin, densely concentrated layers (45). Additionally, in the absence of significant fluid shear (i.e., shear rates < 0.1 s^−1^, which are typically found in the subsurface layers of the ocean), individual elongated cells with a high aspect ratio (> 5) orient themselves with their major axis parallel to gravity within a few minutes and maintain this orientation while sinking (29, 46). Conversely, under strong mixing conditions, cell orientations are random. Cell orientation in pennate diatoms is driven by chloroplast relocation movements, shifting from a position near the nucleus to the valvar ends (and vice-versa), as proposed by Font-Muñoz *et al.,* 2021.

In Figure 3 we demonstrate how the gradual cell reorientation of a suspension of *Pseudo-nitzschia fraudulenta* cells generates anisotropy in the ChlF field emitted by a population, as detected from the two orthogonal red-fluorescence sensors. In the experiment, the initial agitation induces random cell orientations resulting in an isotropic fluorescence as cells progressively orient vertically, the ChlF at orthogonal directions (π/2, π) diverges rapidly. Within the first 60s after the cessation of the agitation exerted in the suspension, 50% of the final anisotropy is already achieved. After a few minutes, the intensity of emitted red light at π/2 is significantly larger than that at π, and this difference steadily increases over time until reaching saturation (Fig. 3A). In contrast, fluorescence emissions at symmetric orientations relative to the vertical reference axis (i.e., at sensor positions 3π/4 and 5π/4) remain similar throughout the experiment, indicating symmetry in the fluorescence field along these angular positions (Fig. 2C).

**Figure 3.**
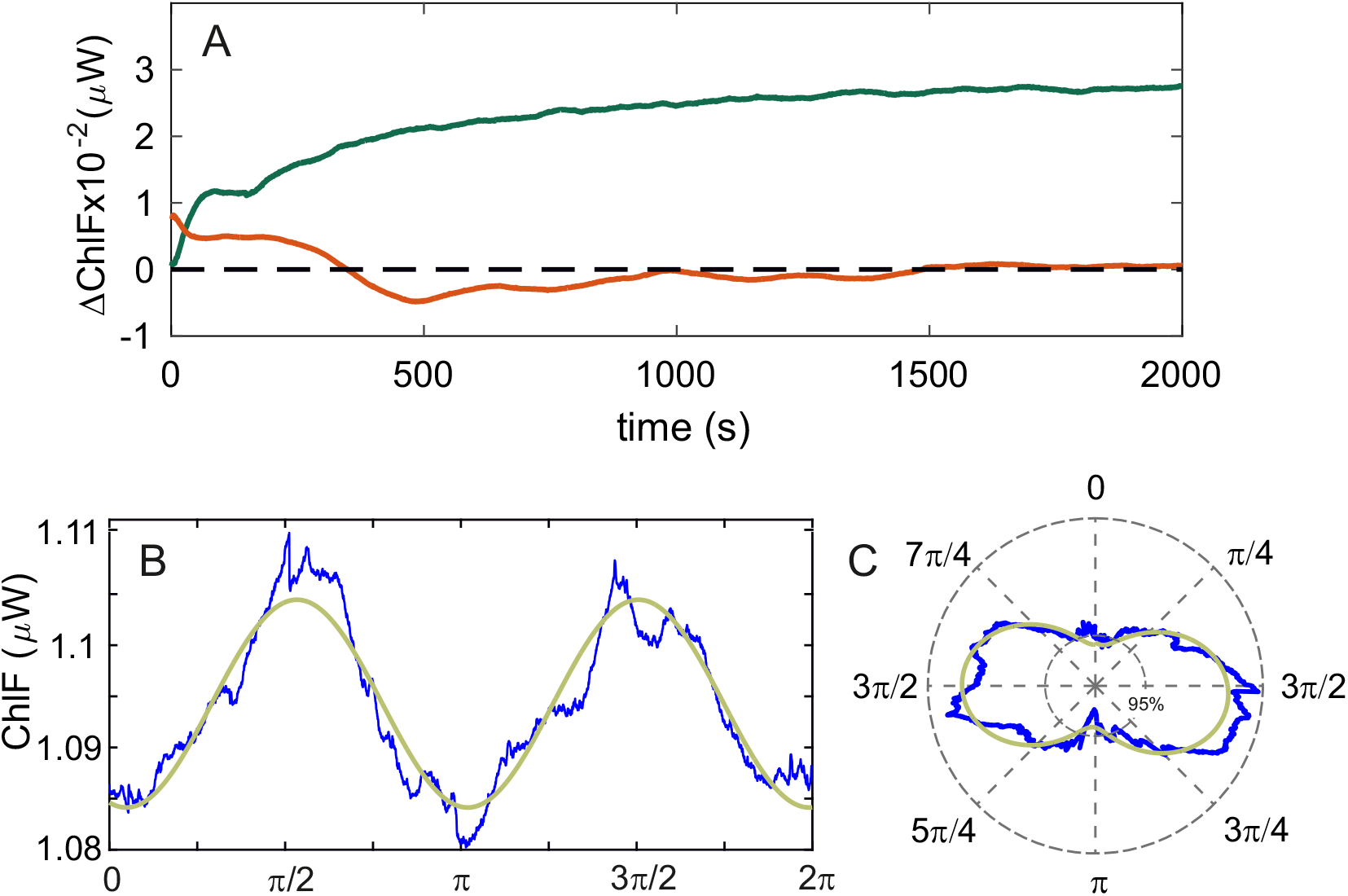
Anisotropy in the radial fluorescence distribution of *Pseudo-nitzschia fraudulenta*. **(A)** Temporal differences in fluorescence intensity (π/2, π, and 3π/4, 5π/4 in green and red respectively) during the experiments. **(B)** Emitted fluorescence from a cell suspension of *Pseudo-nitzschia fraudulenta* as a function of angle (blue line). The green line represents the experimental data, and the red line shows the sinusoidal fit. **(C)** Radial distribution of the fluorescence data shown in panel (B).

To better characterize the ChlF field emitted by a coherently aligned (vertically oriented) population (10^6^ cells L^-1^) of *P. fraudulenta*, we measured the emitted fluorescence at angular intervals of ∼0.01 rad across the entire 2π angular range. Figures 3B and 3C display the resulting intensity field, which is markedly directional. ChlF peaks at π/2 and 3π/2 and is minimum at 0 and π. Since these elongated cells are vertically oriented, we can infer that ChlF is maximized along the apical axis, while the minimum intensity aligns with the transapical axis of cells. This collective generation of ambient ChlF anisotropy (*r*= 4%) is attributed to the directionality in the emission of individual cells and, therefore, should depend on the orientational angular distribution within the population and, hence, on the proportion of cells that are effectively aligned.

To determine whether the observed anisotropy in the collective fluorescence light field emitted by a population of planktonic pennate diatoms is specific to *P. fraudulenta* or represents a more conserved trend, we extended our analysis to two other species of the genus *Pseudo-nitzschia* (*P. delicatissima* and *P. pungens*). As a control, we used *Synechococcus*, a species whose small size (approximately 1 μm) and relatively spherical shape are unlikely to induce preferential orientation, as its alignment is expected to remain largely random. Following the same procedure, we observed that populations of *P. delicatissima* also exhibit marked anisotropy in the radial distribution of ChlF, while *P. pungens* show a somehow weaker signal (r=1.2% and r=0.5% respectively). Conversely, the *Synechococcus* suspension displayed the expected radially isotropic emission (r=0.3%) as shown in Figure 4.

**Figure 4.**
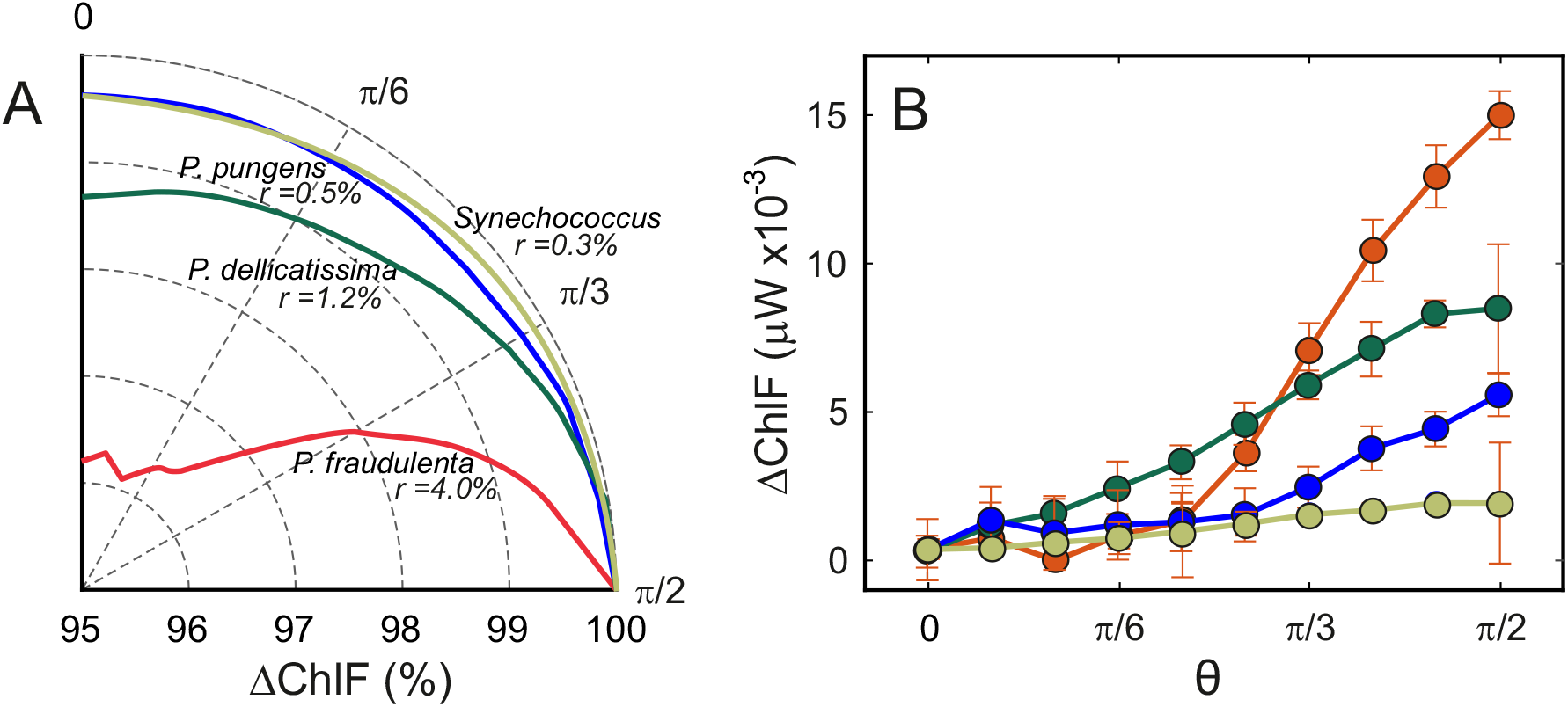
Radial distribution of emitted fluorescence from a cell suspension of different species within the genus *Pseudo-nitzschia* and *Synechococcus* shown as the relative percentual **(A)** and magnitude **(B)** angular dependence.

Notably, the ∼35% difference between maximum and minimum intensity observed in individual cells reduces to ∼5% when measured at a population level in a cell suspension. This discrepancy may arise from several factors, including the integrating effect of the red-light detectors -which capture photons over a defined angular aperture- and the inherent natural variability in the vertical orientation of individual sinking cells (46).

The influence of population-level fluorescence anisotropy on the local light climate is effectively represented using a simplified one-dimensional (1D) model. In this framework, we examine the variations in perceived fluorescence at a fixed vertical distance *z* from a 1D array of uniformly distributed cells (serving as a minimal representation of a thin phytoplankton layer) each emitting anisotropically according to the empirically measured single-cell fluorescence emission pattern (Fig. 2C). Cell orientations within the array follow a distribution parametrized by a scalar value ε ∈ [−1.1], which quantifies deviations from a uniform distribution, as detailed in the Methods section. Figure 5B shows three representative orientation distributions *P*(*θ*) characterized by different values of ε. These orientation distributions reflect conditions expected at different depths within the water column, shaped by prevailing hydrodynamic regimes such as turbulence, shear, or stratification (Fig. 5A). The average normalized fluorescence intensity ⟨*I*_*ε*_(*z*)⟩/⟨*I*_0_(*z*)⟩ - which is, to a first approximation, independent of the vertical distance from the layer - shows a strong dependence with ε as depicted in Figure 5C, with variations of order 10% between cells that are mostly oriented with the maximum intensity orientation and those randomly oriented. These variations are compatible with those measured in collective experiments and highlight that hydrodynamic forces could significantly influence the ambient fluorescence field generated by a diatom population, especially in regions of high concentration of cells.

**Figure 5.**
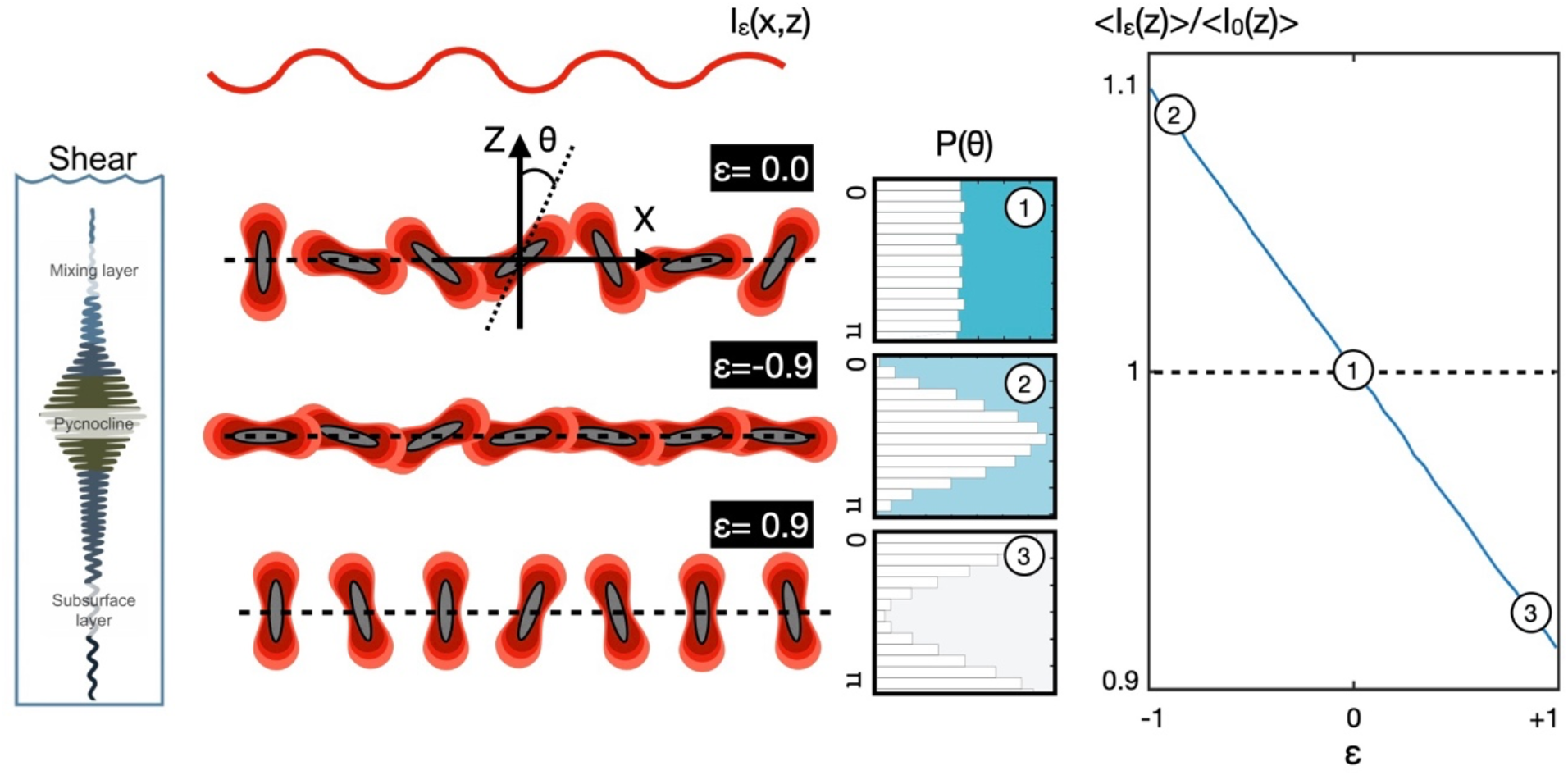
**(A)** Sketch of the variation of the shear stress through the water column. Highest values occure at the pycnocline. **(B)** The effect of the parameter E on the orientation of cells according to the shown orientation distributions *P*(*θ*). *I*_*ε*_(*x, z*) represent the emitted intensity at a certain distance *z* from the plane where the array of cells is located for a given value of ε. **(C)** The dependence of the average emitted intensity for an array of cells normalized with the intensity of randomly oriented cells,*I*_0_(*z*) as a function of E. Specific values for the orientation distributions depicted in (B) are denoted with 1, 2 and 3.

Chain formation, a common trait in certain diatom species, can further enhance anisotropy in the fluorescence field by extending the directional coherence of light emission across multiple aligned cells. In our experiments, while *P. fraudulenta* and *P. delicatissima* populations primarily consist of individual cells, with over 95% of cells found in culture as single cells *P. pungens* exhibits a pronounced tendency to form long apically connected cell chains. Notably, cultures of *P. pungens* contain a substantial mix of both solitary cells and chains exceeding four cells (see Fig. S1). Recent observations indicate that *P. pungens* chains, which exhibit flexible fiber-like behavior during sinking, predominantly orient themselves horizontally, regardless of their initial orientation (29). The coexistence of vertically and horizontally oriented P. pungens chains within cultures likely contributes to a reduction in the overall light anisotropy of the population. Conversely, the differences in ChlF between *P. fraudulenta* and *P. delicatissima* may be attributed to morphological variations. Specifically, *P. fraudulenta* exhibits a larger size, with a major axis of 40μm and a minor axis of 10μm, compared to *P. delicatissima,* which has a major axis of 30μm and a minor axis of 3μm.

Recent studies have demonstrated that the collective orientation of marine microorganisms, influenced by microscale hydrodynamic conditions, affects the optical properties of the water column. For instance, Marcos et al. (2011) showed that changes in the collective orientation of marine phytoplankton blooms can increase optical backscatter by more than 20%. Here, we demonstrate that, in addition to influencing the propagation of sunlight through the water column, collective orientations also influence the emission of autofluorescence at the population level. Fluorescence variations of up to 20% may arise not from changes in cell biomass or physiological status, but solely due to changes in the orientation of individual cells.

The consequences of anisotropy in ChlF extend across multiple spatial scales, influencing processes from the individual cell level to entire populations. At a single-cell level, anisotropy and directionality in the emitted fluorescence field may encode valuable information that other organisms — whether cogeners, competitors, parasites, or predators — can utilize to optimize their positioning and behavior. The anisotropy in the light field emitted by a population of diatoms could have significant ecological implications, particularly when considering how red light attenuates in seawater. According to Lambert–Beer’s law, red fluorescence decays exponentially with distance, implying that the ability to detect a diatom layer through its emitted fluorescence depends heavily on its angular emission pattern. In open ocean waters, for a red wavelength of 660 nm, the attenuation coefficient is approximately *K* = 0.5 m^−1^ (Kirk, 1994). This indicates that even a relatively modest variation in the collective emission field, such as the ∼20% difference predicted between preferential cell orientations, can lead to macroscopic shifts in the distance at which the same fluorescence intensity is perceived (Δr ≈ 0.44 m; see Fig. S2)

From an ecological perspective, such variation could have important implications for biotic interactions. For instance, organisms sensitive to red light, such as certain bacteria containing phytochromes (47), could detect these fluorescence fields to guide processes like phototaxis or foraging (48–50). These microorganisms might be able to detect a diatom layer under one orientation regime but fail to perceive it under another. For example, a layer of vertically oriented diatoms — a configuration commonly found in the ocean due to the relatively fast reorientation timescales of cells (∼600 s) — (46) would emit most of its fluorescence laterally. As a result, such a layer could be detectable when approached horizontally, yet remain virtually invisible when observed from above or below. In this way, the modulation of fluorescence based on collective orientation could influence various ecological interactions, such as predator-prey encounters, competition, or microbial signaling in thin layers or aggregations.

Interestingly, the spatial scale over which such photic signals remain detectable is notably larger than that of chemical cues, which typically operate over much shorter distances in marine environments (Fig. S2). Info-chemicals such as DMSP or quorum sensing molecules often diffuse or become diluted within just a few hundred micrometers to a few millimeters, before being stretched and folded by turbulence (51) and dissipated by diffusion (52, 53). This contrast highlights a key difference between phototaxis and chemotaxis in the ocean: while both rely on environmental gradients, the effective sensory range of light-based cues can extend far beyond that of chemical signals and, in addition, it is faster, cheaper and largely independent of hydrodynamics (in contrast to the stretching and folding of chemical cues in turbulence). This could make anisotropic light emission a particularly efficient channel for long-range ecological communication.

Furthermore, light signals can indirectly induce changes in the metabolism of nearby cells (Wilde and Mullineaux, 2017). At larger scales (i.e., hundreds of meters and beyond), fluorescence emission is also remotely retrievable in the upwelling irradiance spectra, providing valuable insights on a variety of physiological characteristics of marine phytoplankton (Neville and Gower, 1977; Behrenfeld et al., 2009). Several ocean color sensors include fluorescent bands (e.g., MODIS, OLCI on Sentinel-3A/B). However, fluorescence retrieval algorithms commonly assume isotropic emission, an assumption that, as demonstrated in this study, may not always accurately represent the directional characteristics of the fluorescence in the sea.

## CONCLUSION

In the present study, we demonstrate that pennate diatoms emit red autofluorescence in a highly directional pattern, with an anisotropy factor of r=35% at the individual cell level in the case of *Pseudo-nitzschia fraudulenta.* Our findings indicate that this characteristic can eventually scale up to the population level under hydrodynamic conditions that favor cell realignment or chains. This phenomenon may be particularly relevant in environments where dense diatom populations form thin layers, as often observed in upwelling systems and estuaries. Consequences of ChlF anisotropy potentially range from intercell communication and luminous quorum sensing to satellite-based estimation of ocean productivity. However, as revealed by our experiments, the fluorescence directionality at the individual level and spatial scales of cell-to-cell interactions remains a prominent and measurable signal.

ACKNOWLEDGMENTS

J.S.F-M was supported by funding from “Margalida Comas” postdoctoral fellowship PD/018/2020 from Govern de les Illes Balears and Fondo Social Europeo. Funding from PID2019-104232GB-I00 and PID2022-143018NB-I00 grants from the Spanish Ministerio de Ciencia e Innovación (MICINN) and grant RGP007/2024 from the International Human Frontier Science Program Organization (HFSPO) are acknowledged. The present research was carried out within the framework of the activities of the Spanish Government through the “María de Maeztu Centre of Excellence’’ accreditation to IMEDEA (CSIC-UIB) (CEX2021-001198-M).

## METHODS

### Cultures

Cultures were maintained in 80 mL of sterile-filtered oligotrophic seawater amended with nutrients (K/2) at 17°C, under a 12:12 light:dark cycle and a light intensity of 80 μE · m^−2^ · s^−1^, in algal incubators. The strains were initially identified using transmission electron microscopy (TEM) and confirmed by molecular analysis. Genomic DNA was extracted using the DNeasy Plant Mini Kit (cat. no. 69104 and 69106), and polymerase chain reaction (PCR) was performed with the PN PNS-F1 primer: GGA-TCA-TTA- CCA-CAC-CGA-TCC and PSN-R1: CCT-CTT-GCT-TGA-TCT-GAG-ATC-C; (54).

Cultures were diluted once a week to stay in their exponential growing phase with high cell densities (>10^8^ particles · L^−1^).

### Single Cell experiments

*Pseudonitzschia fraudulenta* cells were grown in F/2+Si medium in 12/12 h light/dark cycles. Micromanipulation experiments were carried out in a Zeiss Axiovert 200 microscope as sketched in Figure 1(B). Cells were observed through a Zeiss WI Plan-Apochromat 63x, NA=1.0 objective. A Thorlabs RG645 longpass filter was installed to observe cell autofluorescence. To use the micromanipulation system a custom 3D printed was employed. Cells were held with a Sutter manual microinjection system attached to the Scientifica Patchstar micromanipulation system. To allow the rotation of the cell the microinjector was installed in a high precision motorized rotation mount Thorlabs PRM1/MZ8 electronically controlled through Thorlabs KDC101 K-cube. Micropipettes were pulled with a WPI PUL-1000 pipette puller. In some cases, pipettes were forged with a WPI MF-200 microforge system to adjust the diameter of the tip to the required dimensions of *P. Fraudulenta* cells, namely∼5-8 μm.

Cells were loaded in a 1cmx1cmx2mm PDMS observation chamber with two opposite slits to allow cell micromanipulation and optical fiber illumination. Autofluorescence of the cells was achieved with a Thorlabs LED M470L2 connected to a Thorlabs FT 200 optical fiber which was placed near the cell with a system of mechanical linear stages. Power of the LED was adjusted to yield 100 μmol/(m^2^ s). Brighfield and fluorescence images at a given cell orientation were recorded with a Teledyne Kinetix sCMOS camera once the cell and the optical fiber were placed in focal plane. Experiments were repeated at different orientations performing 5º steps.

### Population level experiments

To study the fluorescence emitted by a population of cells, an experimental system was designed in which a 100 ml culture volume, with a cell density of 10^6^ cells L^-1^ (corresponding roughly to an average intercellular distance of 1 mm assuming a homogeneous distribution), was illuminated with blue light from a Thorlabs M470L2 LED, with an intensity of 100 μmol m^−2^ s^−1^. Powermeters (PM16-130, Thorlabs) were positioned perpendicular to the light source to measure the autofluorescence signal. A Thorlabs RG645 longpass filter was placed before the sensors to block non-red light and allow the exclusive detection of autofluorescence.

The system was mounted on a motorized microrotator (PRM1Z8, Thorlabs), which allowed the sensors to rotate through a whole 2π angular range in the plane perpendicular to the blue light, while the light source remained fixed. This setup enabled the characterization of the autofluorescence field emitted by the phytoplankton population throughout the full rotation. Measurements were taken at intervals of 0.01 rad, ensuring a detailed mapping of the autofluorescence emission

### Model

To gain an understanding of the anisotropic light emission at the population level, we developed a simple 1D model for light emission in a thin phytoplankton layer. We consider a 1D array of uniformally arranged cells distributed in the horizontal at a certain depth z in the water column. The orientation of individual cells in the array is taken from an angular distribution *P*(*θ*) characterized by a single parameter ε that accounts from a first order harmonic deviation from the uniform and that is given by:

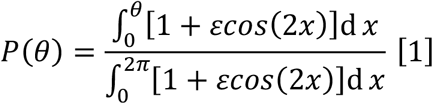

Thus, ε=0 corresponds to a uniform orientation distribution whereas preferential orientation at *θ* = π/2 ∧ π with respect to gravity corresponds with ε = −1 and +1, respectively.

We define *I*_*ε*,_(*x, z*), as the light emitted by cell i with an orientation *θ*, at (*x, z*) as illustrated in Figure 5B. Light emitted by each cell in the array is anisotropic and follows the relative orientation dependence portraited in Fig. 2C with respect to the cell major axis, which we approximante by:

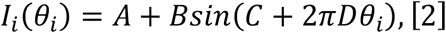

Fitting parameters *A, B, C, D* - obtained by fitting this function to the experimental autofluorescence signal – are A = 80.01 ± 0.15, B = 14.72 ± 0.27, *C* = −1.62 ± 5.6 · 10^(^3 and D = 0.31 ± 1.1 · 10^−5^(Fig.S3).

We average the intensity emitted by the array ⟨*I*_*ε*_(*z*)⟩ at a given vertical distance z, hence neglecting variations in *x*, to yield:

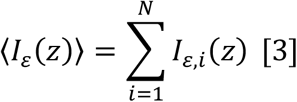

which we normalize with respect to ⟨*I*_0_(*z*)⟩, the average value for the random orientation distribution (ε = 0) as shown in Figure 5C. Since this normalization is performed for the same number of cells, the intensity depicted in figure 5C becomes independent of the number of cells, *N*, for a sufficiently large value of individuals.

## Supplementary material

**Figure S1.**
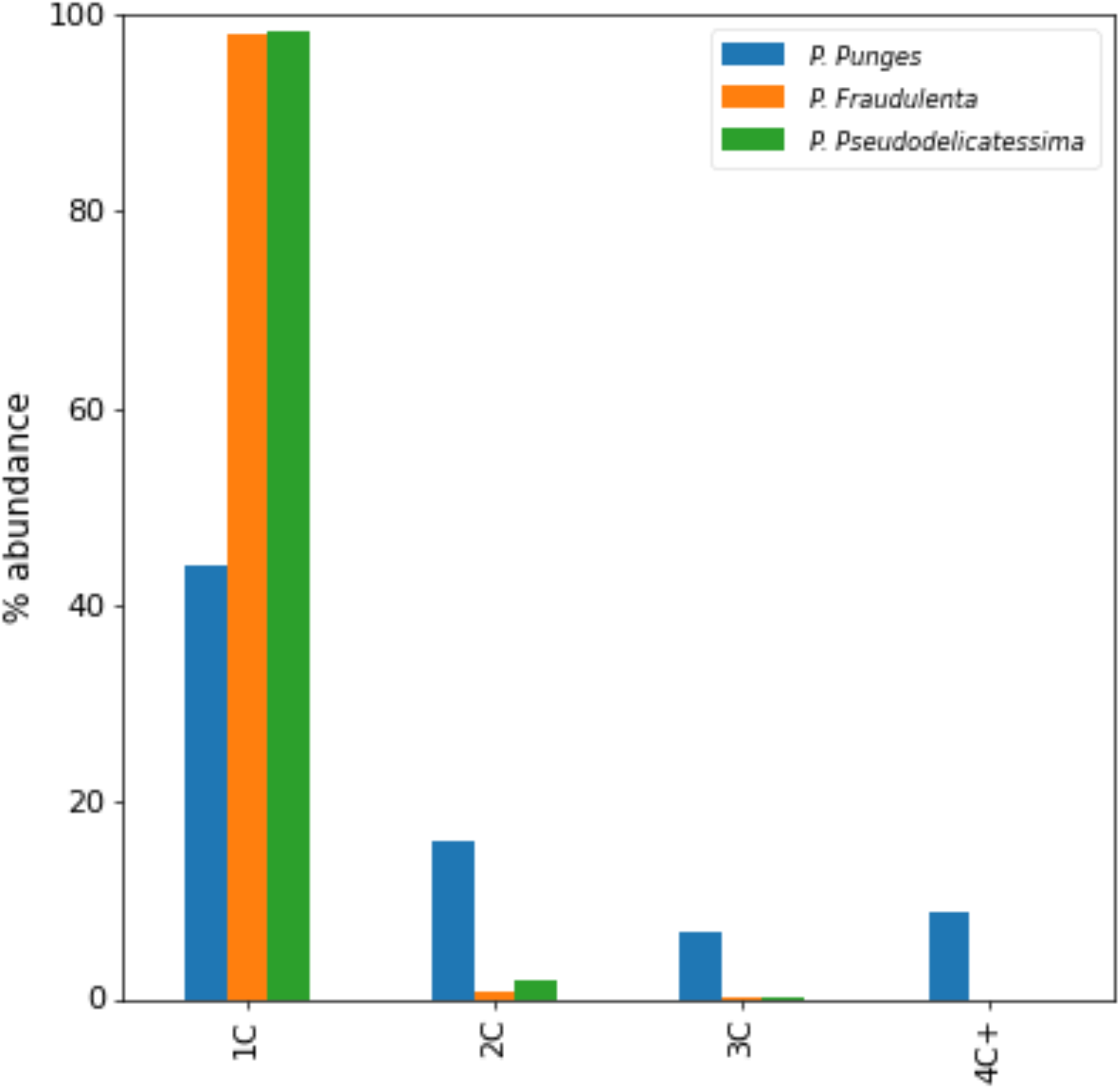
Percentage of individual cells and chains (formed by 2, 3, or more than 4 cells) in the cultures of *Pseudo-nitzschia pungens, Pseudo-nitzschia dellicatissima*, and *Pseudo-nitzschia fraudulenta*.

**Figure S2.**
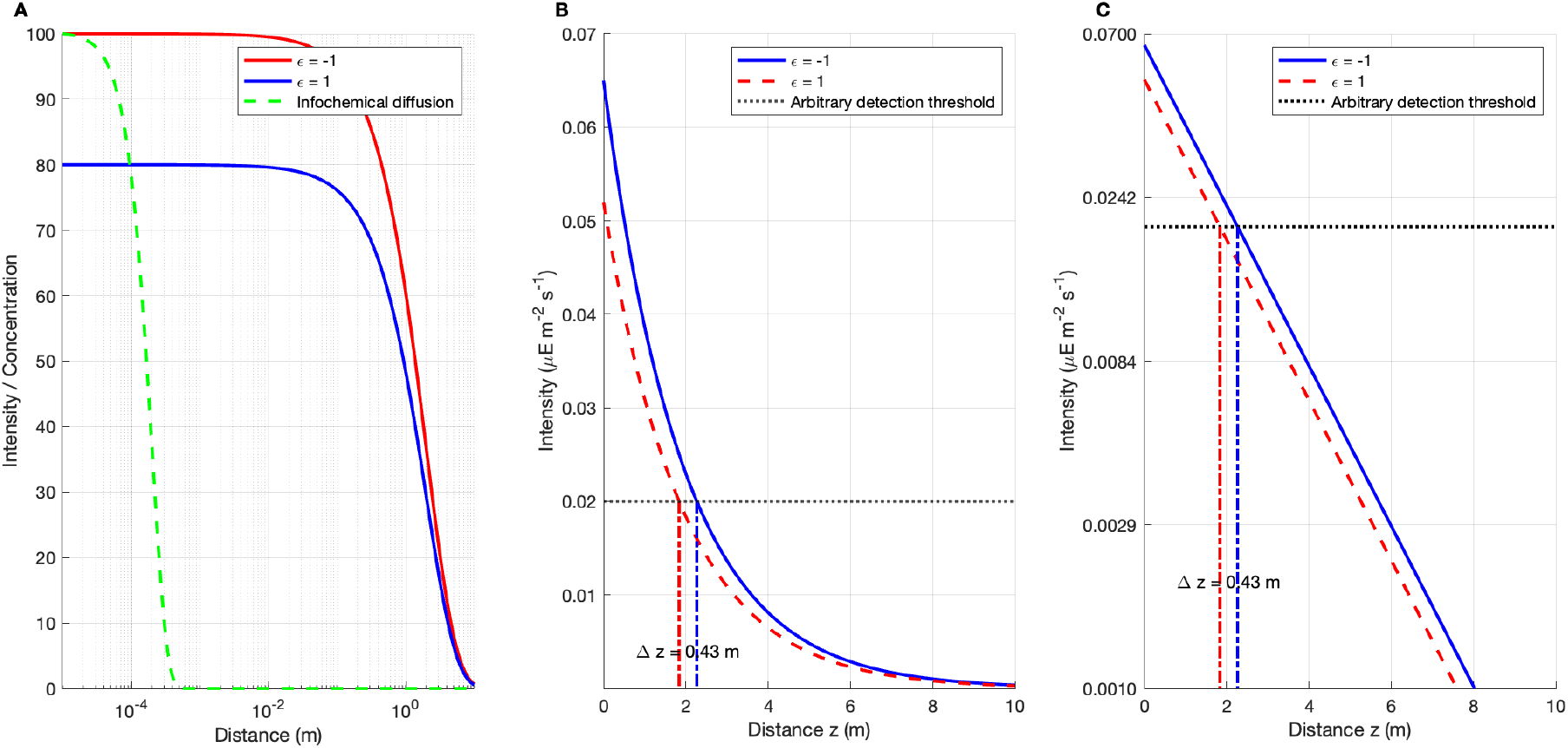
A) Comparison of red light attenuation and infochemical diffusion as a function of distance. Light attenuation follows the Beer–Lambert law, *I*(*z*) = I_0_ · e^-kz.^, with an attenuation coefficient *k = 0.5* m^−1^, representative of red light (∼660 nm) in clear seawater. Two initial intensities are shown: *I*_*0*_ = *100* (ε = −1) and *I*_0_ *= 80* (ε = 1). In contrast, the chemical signal is modeled as the diffusion of a molecule from a point source in a three-dimensional medium, with concentration 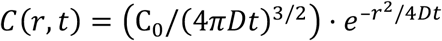, where *D = 10^*−9*^* m^2^/s. This leads to a much sharper spatial decay (submillimeter range), emphasizing the difference between the spatial scales of phototactic and chemotactic cues. B) Attenuation curves of fluorescent light for both ϵ values. The initial light intensities used correspond to measured fluorescence emission from a population with a density of *10^6^* cells/L. A horizontal dashed line represents an arbitrary detection threshold, and vertical lines indicate the detection distances. The horizontal distance between them quantifies the sensitivity range difference (*Δz = 0.43* m). C) Same data as in panel B, shown in log-transformed intensity values. The two curves are parallel, illustrating that the difference in detection range (*Δz*) remains constant across any arbitrary threshold.

**Figure S3.**
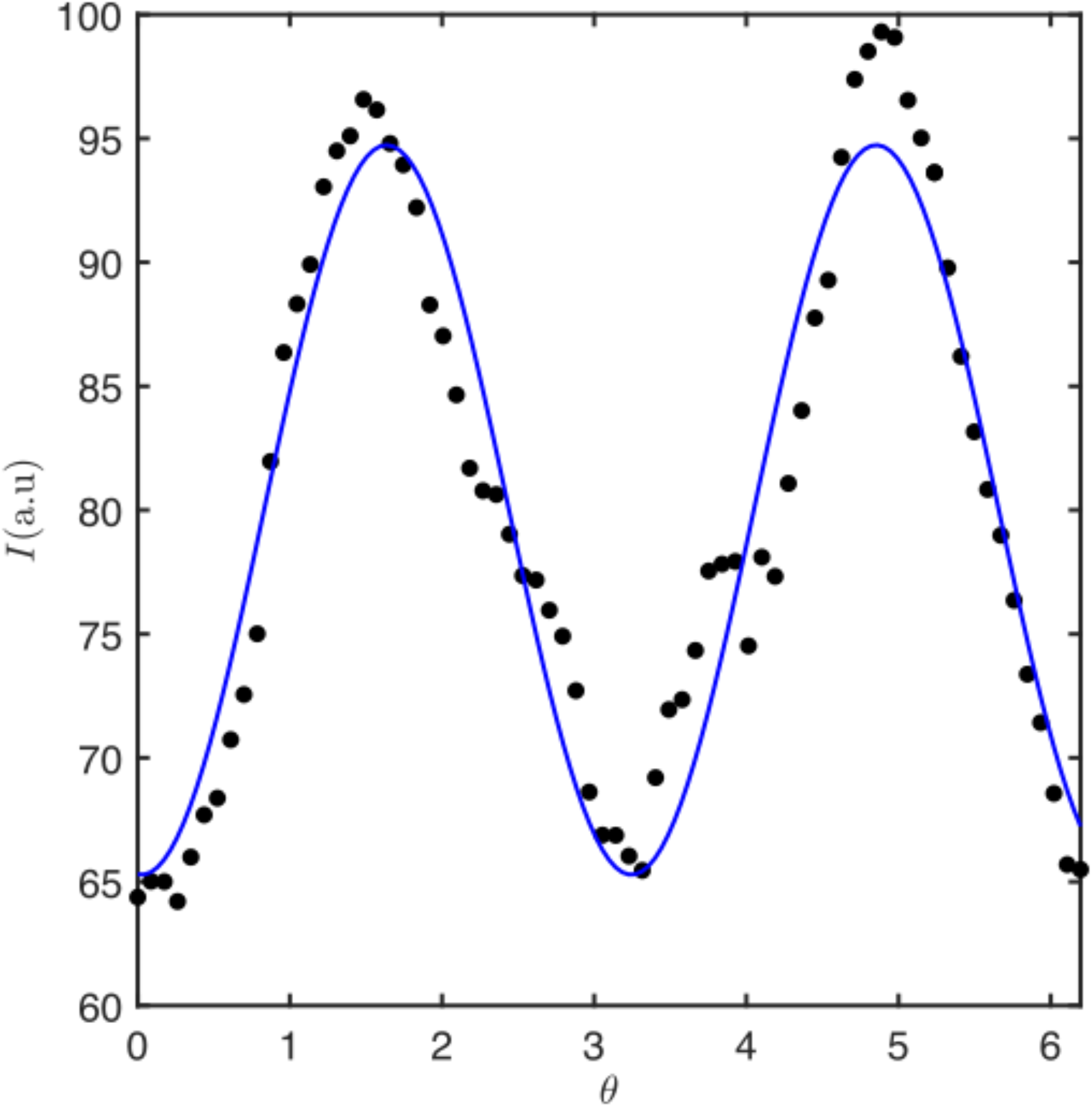
Black dots represent experimental data of fluorescence emission measured at the single-cell level. The blue line shows a sinusoidal fit to these experimental data.

## REFERENCES

1. Hink MA, et al. (2000) Structural dynamics of green fluorescent protein alone and fused with a single chain Fv protein. J Biol Chem 275(23):17556–17560.

2. Marshall J, Johnsen S (2017) Fluorescence as a means of colour signal enhancement. Philos Trans R Soc B Biol Sci 372(1724). doi:10.1098/rstb.2016.0335.

3. Liu X, et al. (2021) Chlorophyll fluorescence as a light signal enhances iron uptake by the marine diatom Phaeodactylum tricornutum under high-cell density conditions. BMC Biol 19(1):1–15.

4. Gerlach T, Sprenger D, Michiels NK (2014) Fairy wrasses perceive and respond to their deep red fluorescent Coloration. Proc R Soc B Biol Sci 281(1787). doi:10.1098/rspb.2014.0787.

5. Gruber DF, Sparks JS (2015) First observation of fluorescence in marine turtles. Am Museum Novit 2015(3845):1–8.

6. García-Plazaola JI, et al. (2015) Autofluorescence: biological functions and technical applications. Plant Sci 236:136–145.

7. Macel M-L, et al. (2020) Sea as a color palette: the ecology and evolution of fluorescence. Zool Lett 6(1):9.

8. Poding LH, Jägers P, Herlitze S, Huhn M (2024) Diversity and function of fluorescent molecules in marine animals. Biol Rev. doi:10.1111/brv.13072.

9. Croce AC (2021) Light and autofluorescence, multitasking features in living organisms. Photochem 1(2):67–124.

10. Mondal PP, Diaspro A, Mondal PP, Diaspro A (2014) Basics of fluorescence and photophysics. Fundam Fluoresc Microsc Explor Life with Light:111–134.

11. Hamchand R, et al. (2021) Red Fluorescence of European Hedgehog (Erinaceus europaeus) Spines Results from Free-Base Porphyrins of Potential Microbial Origin. J Chem Ecol 47(6):588–596.

12. Arnold KE, Owens IPF, Marshall NJ (2002) Fluorescent signaling in parrots. Science (80-) 295(5552):92.

13. Meadows MG, et al. (2014) Red fluorescence increases with depth in reef fishes, supporting a visual function, not UV protection. Proc R Soc B Biol Sci 281(1790). doi:10.1098/rspb.2014.1211.

14. Jerlov NG (1976) Marine optics (Elsevier).

15. Kirk JTO (1994) Light and photosynthesis in aquatic ecosystems (Cambridge university press).

16. Lagorio MG, Cordon GB, Iriel A (2015) Reviewing the relevance of fluorescence in biological systems. Photochem Photobiol Sci 14(9):1538–1559.

17. Butler WL (1978) Energy distribution in the photochemical apparatus of photosynthesis. Annu Rev Plant Physiol 29(1):345–378.

18. Gorbunov MY, Falkowski PG (2022) Using chlorophyll fluorescence to determine the fate of photons absorbed by phytoplankton in the world’s oceans. Ann Rev Mar Sci 14(1):213–238.

19. Falkowski P, Kiefer DA (1985) Chlorophyll a fluorescence in phytoplankton: relationship to photosynthesis and biomass. J Plankton Res 7(5):715–731.

20. Chamberlin WS, Booth CR, Kieffer DA, Morrow JH, Murphy RC (1990) Evidence for a simple relationship between natural fluorescence, photosynthesis and chlorophyll in the sea. Deep Sea Res Part A Oceanogr Res Pap 37(6):951– 973.

21. Chamberlin S, Marra J (1992) Estimation of photosynthetic rate from measurements of natural fluorescence: analysis of the effects of light and temperature. Deep Sea Res Part A Oceanogr Res Pap 39(10):1695–1706.

22. Kiefer DA, Reynolds RA (1992) Advances in understanding phytoplankton fluorescence and photosynthesis. Prim Product Biogeochem cycles sea:155–174.

23. Stegmann PM, Lewis MR, Davis CO, Cullen JJ (1992) Primary production estimates from recordings of solar-stimulated fluorescence in the equatorial Pacific at 150° W. J Geophys Res Ocean 97(C1):627–638.

24. Walker D (1987) The use of the oxygen electrode and fluorescence probes in simple measurements of photosynthesis (Citeseer).

25. Falkowski PG, Raven JA (2013) Aquatic photosynthesis (Princeton University Press).

26. Cullen JJ, Ciotti AM, Davis RF, Neale PJ (1997) Relationship between near-surface chlorophyll and solar-stimulated fluorescence: biological effects. Ocean Optics XIII (SPIE), pp 272–277.

27. Behrenfeld MJ, et al. (2009) Satellite-detected fluorescence reveals global physiology of ocean phytoplankton. Biogeosciences 6(5):779–794.

28. Font-Muñoz JS, Sourisseau M, Cohen-Sánchez A, Tuval I, Basterretxea G (2021) Pelagic diatoms communicate through synchronized beacon natural fluorescence signaling. Sci Adv 7(51):eabj5230.

29. Sourisseau M, et al. (2024) Sinking rates, orientation and behavior of pennates diatoms. J Phycol.

30. Font-Munoz JS, et al. (2024) Phytochromes Enable Social Behavior in Marine Diatoms. bioRxiv:2024.09.18.613651.

31. Long JD, Smalley GW, Barsby T, Anderson JT, Hay ME (2007) Chemical cues induce consumer-specific defenses in a bloom-forming marine phytoplankton. Proc Natl Acad Sci 104(25):10512–10517.

32. Vardi A (2008) Cell signaling in marine diatoms. Commun Integr Biol 1(2):134– 136.

33. Armbrust EV (2009) The life of diatoms in the world’s oceans. Nature 459(7244):185.

34. Bowler C, Vardi A, Allen AE (2010) Oceanographic and biogeochemical insights from diatom genomes. Ann Rev Mar Sci 2:333–365.

35. Trainer VL, et al. (2012) Pseudo-nitzschia physiological ecology, phylogeny, toxicity, monitoring and impacts on ecosystem health. Harmful Algae 14:271– 300.

36. Mcfarland M, Nayak AR, Stockley N, Twardowski M (2020) Enhanced Light Absorption by Horizontally Oriented Diatom Colonies. 7(July):1–10.

37. D’Mello Y, et al. (2022) Solar energy harvesting mechanisms of the frustules of Nitzschia filiformis diatoms. Opt Mater Express 12(12):4665.

38. Marcos, et al. (2011) Microbial alignment in flow changes ocean light climate. Proc Natl Acad Sci 108(10):3860–3864.

39. Mann DG (1996) Chloroplast morphology, movements and inheritance in diatoms. Cytol Genet Mol Biol algae.

40. Fu W, et al. (2022) Diatom morphology and adaptation: Current progress and potentials for sustainable development. Sustain Horizons 2:100015.

41. Drum RW (1963) The cytoplasmic fine structure of the diatom, Nitzschia palea. J Cell Biol 18(2):429–440.

42. Goessling JW, et al. (2018) Structure-based optics of centric diatom frustules: modulation of the in vivo light field for efficient diatom photosynthesis. New Phytol 219(1):122–134.

43. Goessling JW, Yanyan S, Kühl M, Ellegaard M (2021) Frustule photonics and light harvesting strategies in diatoms. Diatom Morphog:269–300.

44. De Tommasi E (2016) Light Manipulation by Single Cells: The Case of Diatoms. J Spectrosc 2016. doi:10.1155/2016/2490128.

45. Nayak AR, McFarland MN, Sullivan JM, Twardowski MS (2018) Evidence for ubiquitous preferential particle orientation in representative oceanic shear flows. Limnol Oceanogr 63(1):122–143.

46. Font-Muñoz JS, et al. (2019) Collective sinking promotes selective cell pairing in planktonic pennate diatoms. Proc Natl Acad Sci U S A 116(32):15997–16002.

47. Falciatore A, Bowler C (2005) The evolution and function of blue and red light photoreceptors. Curr Top Dev Biol 68:317–350.

48. Jékely G (2009) Evolution of phototaxis. Philos Trans R Soc B Biol Sci 364(1531):2795–2808.

49. Armitage JP, Hellingwerf KJ (2003) Light-induced behavioral responses (phototaxis’) in prokaryotes. Photosynth Res 76:145–155.

50. Davis SJ, Vener A V, Vierstra RD (1999) Bacteriophytochromes: phytochrome-like photoreceptors from nonphotosynthetic eubacteria. Science (80-) 286(5449):2517–2520.

51. Batchelor GK (1959) Small-scale variation of convected quantities like temperature in turbulent fluid Part 1. General discussion and the case of small conductivity. J Fluid Mech 5(1):113–133.

52. Stocker R, Seymour JR (2012) Ecology and physics of bacterial chemotaxis in the ocean. Microbiol Mol Biol Rev 76(4):792–812.

53. Raina J-B, Fernandez V, Lambert B, Stocker R, Seymour JR (2019) The role of microbial motility and chemotaxis in symbiosis. Nat Rev Microbiol 17(5):284– 294.

54. Noyer C, Abot A, Trouilh L, Leberre VA, Dreanno C (2015) Phytochip: development of a DNA-microarray for rapid and accurate identification of Pseudo-nitzschia spp and other harmful algal species. J Microbiol Methods 112:55–66.

